# The intrinsically disordered N-terminal region of eIF5B nucleates RNA-driven phase separation

**DOI:** 10.64898/2026.11.02.763741

**Authors:** Brandt Bodley-Gomes, Michael T. Marr

**Author notes:** For correspondence: Michael T. Marr II.

## Abstract

Recently, we found that eukaryotic Initiation Factor 5B (eIF5B) forms cytoplasmic puncta under conditions of cellular stress and that the puncta formation drives an increase in alternative translation initiation. The formation of these puncta depends on the intrinsically disordered N-terminal region of eIF5B (eIF5B-NTR). To address the driving force behind this observation, we reconstituted a purified, binary, in vitro system to study the phase separation potential of the eIF5B-NTR. Utilizing this system, we show that eIF5B-NTR forms RNA-driven condensates at physiological concentrations. Furthermore, we defined the properties of the eIF5B condensates and the amino acid sequence requirements of this polypeptide to form complex liquid-liquid condensates. Truncation mapping identified a central arginine-rich core (residues ∼200-400) that drives condensation but forms nearly static droplets in the absence of the rest of the polypeptide. Finally, we tested this model in cells to determine if the in vitro observations apply to the complex cellular environment.

## Introduction

The cytoplasm has traditionally been viewed as a homogeneous, fluid matrix in which membrane-bound organelles are suspended. However, advances in molecular tagging and high-resolution imaging have challenged this view and revealed a far more structured intracellular environment. During cellular events, including replication and differentiation, or in response to cellular stress, proteins and nucleic acids can become enriched into discrete puncta or granules across nearly all cellular compartments (1–3). These assemblies are commonly referred to as membraneless organelles (MLOs) or biomolecular condensates and are dynamic structures that arise through networks of transient and multivalent intermolecular interactions (4).

Considerable effort has been devoted to understanding the molecular determinants governing biomolecular condensation (5–8). The systematic characterization of such proteins has revealed several functional classes defined by their molecular requirements and sequence features. Many of these model proteins interact with RNA or DNA and contain intrinsically disordered regions (IDRs) enriched in low-complexity sequences (2). Within these proteins, several classes of interactions have been implicated in the formation of condensates, including π-π interactions, π-cation interactions, patterned electrostatic interactions, and multivalent modular binding interfaces (6). For example, the phase separation of the Rubisco linker protein (PYCO1) is proposed to depend primarily on aromatic π-π interactions, whereas π-cation interactions contribute to condensation of the RNA-binding protein Fused in Sarcoma (FUS) (9, 10). Patterned electrostatic interactions have been described for the DEAD-box helicase DDX4, while multivalent domain-ligand interactions drive assembly of systems such as the N-WASP-NCK-phosphorylated nephrin signaling network through SH2 and SH3 domain interactions (11–13).

Phase-separating proteins can be further divided by their dependence on scaffolding partners. Some proteins are capable of independently undergoing simple condensation. For example, the P-granule protein LAF-1 from *Caenorhabditis elegans* can form condensates in vitro absent any binding partners (14). In contrast, many phase-separating proteins require the addition of partners to undergo complex coacervation. A well-studied example can be seen in the RNA-dependent assembly of condensates mediated by the human stress granule protein G3BP1 (15). In addition to sequence-encoded features, post-translational modification of scaffold proteins can modulate condensate formation and material properties, providing an additional regulatory layer (16).

Within cells, membraneless organelles may form constitutively or dynamically in response to physiological cues. Constitutive condensates include structures such as the nucleolus, where the transcription, processing, and maturation of ribosomes are clustered (17–21). In contrast, inducible condensates arise in response to cellular stress or signaling events; for example, stress granules assemble following activation of the integrated stress response and phosphorylation of eIF2α (22, 23). Beyond maintaining cellular homeostasis, phase separation has emerged as an important mechanism in cellular differentiation and developmental biology. The formation of P-granules during *C. elegans* embryogenesis, mediated in part by LAF-1, illustrates how condensates can organize developmental processes (14, 24). Dysregulation of biomolecular condensation is increasingly associated with disease, particularly neurodegenerative disorders, although its roles in developmental, oncogenic, and metabolic pathologies are also becoming apparent (25–27).

In previous work, our laboratory identified the eukaryotic Initiation Factor 5B (eIF5B) as a protein capable of forming biomolecular condensates in the cytoplasm in response to cellular stress (28). Notably, eIF5B condensates appeared distinct from previously characterized cytoplasmic stress-responsive granules and gave rise to an increase in translation output for internal-ribosomal entry site (IRES) mediated transcripts. The distinct localization of eIF5B is consistent with previous proximity-labeling and mass-spectrometry studies that have catalogued protein constituents of stress granules (SGs) and processing bodies (PBs) (29–31). Among canonical components of the translation initiation machinery, eIF5B uniquely fails to localize to either of these established granule types in human cells. Together, these observations led us to suggest that eIF5B nucleates a previously uncharacterized cytoplasmic condensate associated with translational control. Curiously, the heightened expression of eIF5B is also strongly correlated with negative outcomes across a spectrum of cancer types (32, 33). The poor prognosis associated with increased eIF5B expression is thought to be modulated through the non-canonical translation of pro-survival and immune-evasion proteins under conditions of cellular stress (33, 34). A clear link between both modes of translational regulation remains unexplored.

Structurally, eIF5B can be divided into two domains of comparable size. The C-terminal domain (CTD) forms a highly conserved, well-folded catalytic core. Structural and biochemical studies have shown that the CTD is sufficient to mediate the canonical functions of eIF5B in translation initiation, including stabilizing initiator tRNA on the ribosome and promoting large ribosomal subunit joining (35, 36). In contrast, the N-terminal region (NTR) remains poorly characterized. Sequence analyses predict that the NTR is largely intrinsically disordered and enriched in low-complexity segments, consisting primarily of glutamate- and lysine-rich stretches arranged in alternating patches (Fig. 1A) (37).

**Figure 1.**
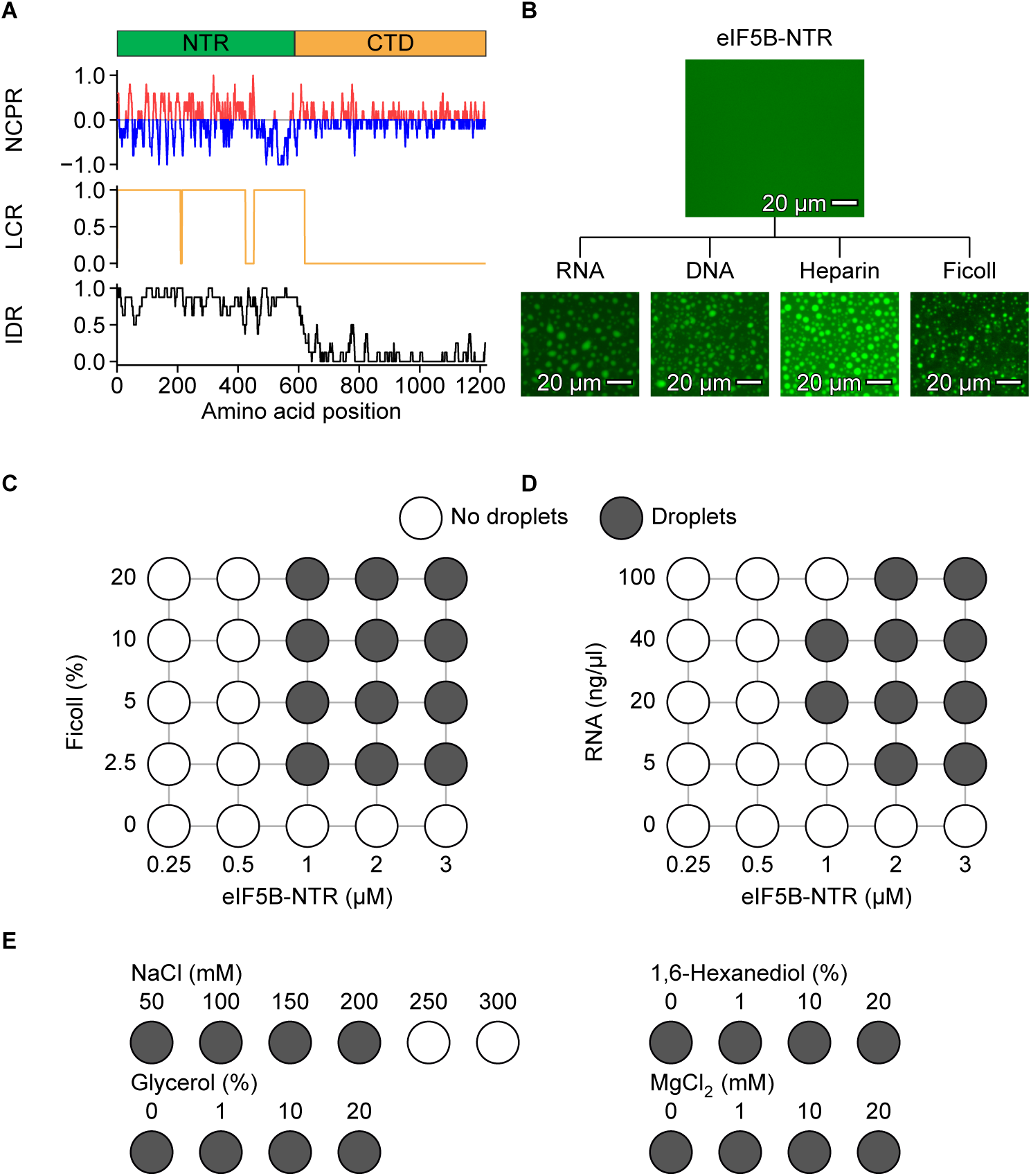
The eIF5B N-terminal region is intrinsically disordered, highly charged and undergoes phase separation in vitro. **(A)** Domain organization of eIF5B (NTR, residues 1-587; CTD, residues 588-1220) with predicted sequence properties along the full-length protein. From top to bottom, the plots show net charge per residue (NCPR; positive values red, negative values blue), low-complexity region assignment (LCR) and predicted intrinsic disorder (IDR). Properties were calculated with MolPhase (37). **(B)** Representative fluorescence micrographs of purified eIF5B-NTR (3.3 µM), with 2% mNG-eIF5B-NTR tracer, shown alone and after addition of RNA (20ng/µl), DNA (20ng/µl), heparin (0.02ng/µl), or Ficoll (5% w/v). Scale bars, 50 µm. **(C)** Phase diagram of eIF5B-NTR as a function of protein (0.25-3 µM) and Ficoll (0-20% w/v) concentration. **(D)** Phase diagram of eIF5B-NTR as a function of protein (0.25-3 µM) and RNA (0-80 ng/µl) concentration. **(E)** Condensate formation by eIF5B-NTR (3.3 µM) with RNA (20 ng/µl) in the presence of increasing NaCl (50-300 mM), 1,6-hexanediol (0-20% v/v), glycerol (0-20% v/v) or MgCl_2_ (0-20 mM). In **C-E**, grey circles indicate conditions with droplets, and white circles indicate conditions with no droplets. A condition was scored as positive when at least 10 droplets were visible in at least two of three independent replicates.

Overexpression of eIF5B-NTR alone in mammalian cells is sufficient to induce formation of full-length eIF5B-containing cytoplasmic granules (28). Despite this observation, the molecular basis of eIF5B condensation and the mechanisms governing its assembly remain unknown. In particular, it is unclear whether the eIF5B-NTR itself possesses intrinsic phase-separating capacity or whether its recruitment into the stress-induced condensate depends on interactions with additional cellular components. Resolving this question is important as it may reveal a previously unrecognized property of the translation machinery associated with translational regulation under stress conditions.

Here, we sought to determine whether the intrinsically disordered N-terminal region of eIF5B is sufficient for biomolecular phase separation. To address this question, we developed a simplified in vitro model system to investigate the condensation behavior of the eIF5B NTR and its interaction partners. Using this system, we find that the eIF5B NTR is capable of phase separation in the presence of crowding agents or polyanions and that these polyanions can modulate the characteristics of the condensates. In addition, we characterized the biochemical interactions that modulate the properties of the condensates in vitro and in cells.

## Results

### eIF5B-NTR is sufficient for phase separation

To test whether the amino-terminal, intrinsically disordered region of eIF5B (eIF5B-NTR; residues 1-587) is capable of phase separation, we expressed and purified the eIF5B-NTR protein from *E. coli* (Fig. S1A, B). Droplet formation was assayed under near-physiological buffer conditions. For ease of visualization, all samples also contained a 2% tracer generated by expressing and purifying from *E. coli* an amino-terminal fusion of monomeric NeonGreen (mNG) to the eIF5B-NTR. In the absence of other components, eIF5B-NTR remains uniformly dispersed (Fig. 1B).

To define the conditions under which phase separation occurs, we performed two-dimensional titrations varying protein and Ficoll concentrations. Ficoll is a synthetic sucrose polymer that lacks ionic charge and acts as a crowding agent often used to test phase separation (15, 38, 39). The eIF5B-NTR protein forms fluorescent droplets of varying size with a range of Ficoll concentrations, 2.5-20% (Fig. 1B, C). The ability to trigger condensation primarily through molecular crowding highlights that the interactions between eIF5B-NTR monomers are fully sufficient for condensation upon crowding.

### eIF5B-NTR undergoes RNA-dependent phase separation in the absence of crowding reagents

In the absence of crowding reagents, eIF5B-NTR remains uniformly disperse in solution. We wondered if there was a biologically relevant component that could facilitate eIF5B-NTR’s condensation. Previously, we have shown that the eIF5B cellular condensates contain RNA, so we tested if RNA was capable of inducing condensation (28). Addition of a 300-nucleotide transcribed RNA induced the formation of micron-scale droplets. Protein titration under these conditions found a saturation concentration (C_sat_) of approximately 1 µM protein with 20 ng/µl RNA. Droplet formation was reversible upon RNase treatment or increased ionic strength (0.25 M NaCl), indicating that phase separation is RNA-dependent and likely mediated by electrostatic interactions (Fig. S1C). Similar droplet formation was observed upon addition of alternative polyanions, including DNA and heparin (Fig. 1B), suggesting that eIF5B-NTR undergoes phase separation with a range of negatively charged polymers and is nucleated through electrostatic interactions with a multivalent anion.

To probe the basis of phase separation, we assessed the sensitivity of droplets to ionic strength and small molecule perturbants (Fig. 1E). Increasing monovalent ion concentrations with sodium chloride (NaCl) led to dissolution of condensates above 250 mM, consistent with a dominant role for electrostatic interactions. Supporting this model, treatment with 1,6-hexanediol or glycerol up to 20% had minimal effect on droplet integrity, indicating that hydrophobic interactions are unlikely to be a primary driving force of phase separation. Together, these results demonstrate that eIF5B-NTR is capable of phase separation and undergoes RNA-dependent, electrostatically driven liquid-liquid phase separation under physiologically relevant conditions.

### RNA-driven condensates are fluid with material properties modulated by the composition and quantity of RNA

In establishing our RNA-driven, eIF5B-NTR condensates, we observed that droplets undergo spontaneous fusion events, consistent with liquid-liquid phase separation (Fig. 2A). To characterize the internal dynamics of eIF5B-NTR condensates, we performed fluorescence recovery after photobleaching (FRAP) using 3.3 µM eIF5B-NTR, 2% fluorescently labeled protein, and 40 ng/µl RNA. Altering the fraction of labelled to unlabeled protein did not measurably alter condensate properties up to 50% incorporation, validating its use for quantitative measurements (Fig. S2). FRAP analysis revealed rapid fluorescence recovery (t_1/2_ = 3.5 s at 40 ng/µl RNA) and a negligible immobile fraction (<2%), indicating that eIF5B-NTR condensates exhibit liquid-like dynamics (Fig. 2A, B). Increasing RNA concentration promoted droplet growth and abundance while also enhancing internal dynamics, as reflected by increased relative fluidity (Fig. 2C). This result illustrates that RNA not only nucleates condensate formation but also tunes its material properties.

**Figure 2.**
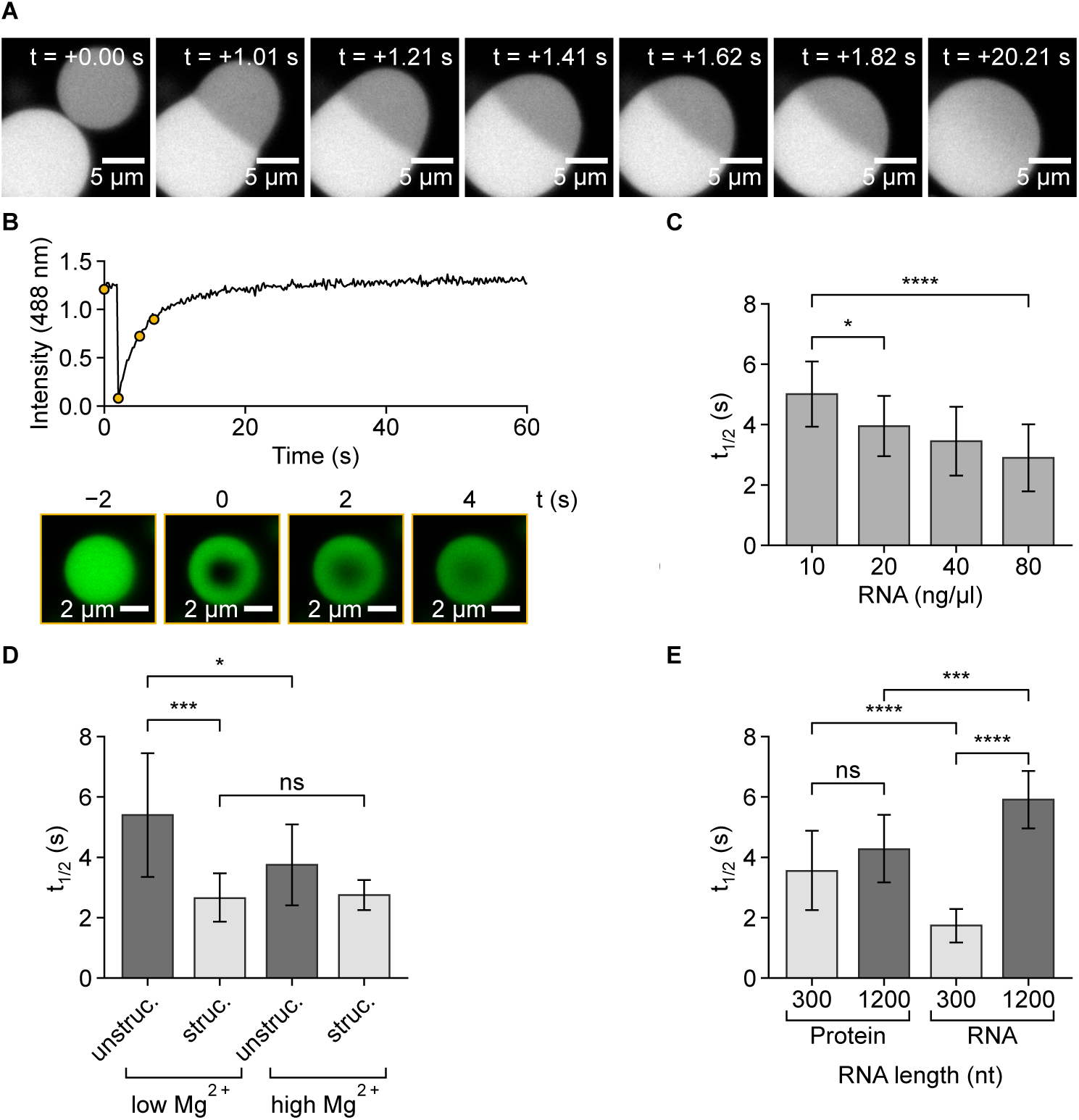
RNA concentration, structure and length tune the internal dynamics of eIF5B-NTR condensates. **(A)** Consecutive frames spanning the fusion of two droplets, imaged at ∼0.2 s intervals and at a final time of 20.21 s. One droplet was partially bleached prior to the time course. Droplets fuse, mix and display rapid shape relaxation. Scale bars, 5 µm **(B)** Representative FRAP recovery trace with confocal images for eIF5B-NTR droplet (2% mNG-eIF5B-NTR) with 40 ng/µl RNA. Confocal images at 2 s intervals spanning prebleach and 4 s post bleach. Yellow dots on recovery curve indicate the time points of bleached images with times are relative to the bleach. Scale bars, 2 µm. **(C)** Recovery time (t_1/2_) of condensates formed with 10-80 ng/µl RNA. **(D)** Recovery time of condensates formed with unstructured or structured model RNA at low (2 mM) or high (10 mM) MgCl_2_. **(E)** Two-color FRAP of protein and fluorescently labeled RNA within condensates assembled with 300- or 1200-nucleotide RNA; x-axis labels give RNA length and the component measured. Data in C-E are mean ± SD of 15 droplets from three independent replicates. Unpaired t test: ns, p ≥ 0.05; *, p < 0.05; ***, p < 0.001; ****, p < 0.0001.

To determine whether RNA structure influences phase behavior, we compared structured and unstructured RNAs of defined lengths, 600 or 1200 nucleotides. To generate a structured RNA, we utilized the well-folded and characterized hepatitis C viral IRES (HCV). The unstructured reference was generated using the beta-globin UTR (BG). Both sequences were supplemented with additional sequence from the open reading frame of Gaussia luciferase (gLuc) to reach the desired length. Of note, both transcripts maintained very similar levels of GC content (55.5 vs 57.3%). Data from FRAP utilizing the 600-nucleotide RNAs (BG and HCV) is shown in Figure 2D. The 1200-nucleotide RNAs produced comparable results (Fig. S2B). Across all samples, RNA structure had minimal impact on phase separation propensity, with all RNAs producing a 1uM C_sat_ when measured with 40 ng/µl RNA. However, more structured RNAs consistently produced more dynamic condensates, with nearly two-fold faster FRAP recovery rates relative to unstructured RNAs of the same length (Fig. 2D). To further test the role of RNA compaction, we increased Mg^2+^ concentration from 2mM to 10 mM, which has previously been shown to promote RNA folding (40, 41). Elevated Mg^2+^ selectively increased the fluidity of condensates formed with unstructured RNA, while having minimal effect on already structured RNA. These results suggest that RNA compaction enhances protein mobility within condensates.

### Protein and RNA exhibit distinct mobility within condensates

Given the observed dependence of condensate properties on RNA character, we next asked whether protein and RNA exhibit similar dynamics within the condensed phase. To address this, we used in vitro transcription to generate RNAs from the previously described BG-gluc transcript at 300 and 1200 nucleotide lengths. We labeled the RNAs at the 3’ end using hydrazide CF^®^594. By assembling a system of fluorescently labelled RNA and eIF5B-NTR with 2% mNG-eIF5B-NTR, we performed two-color FRAP to independently measure the mobility of both components (Fig. 2E). For the smaller 300-nucleotide RNA, the RNA component recovered approximately twice as rapidly as the protein, indicating higher mobility despite its larger molecular size. When we switched to using a 1200-nucleotide RNA, the fluidity of RNA decreased, while the change in protein fluidity was not statistically significant. The threefold decrease in RNA fluidity, in contrast to invariant protein mobility, indicates that the diffusion of protein and RNA within condensates is decoupled.

### A central core region of eIF5B-NTR is required for phase separation

The eIF5B-NTR is predicted to be an intrinsically disordered region with low sequence complexity that is enriched in charged residues. These charged residues exist in alternating patches throughout the NTR with a particularly pronounced acidic stretch at the C-terminal end of this region (Fig. 1A). To identify amino acid sequence elements required for phase separation, we expressed and purified overlapping truncations spanning the NTR and assessed their ability to form RNA-induced droplets in vitro (Fig. 3A). This analysis revealed a central region (residues ∼200-400) that is required for phase separation, whereas N-and C-terminal regions alone are insufficient, even at elevated concentrations (Fig. 3B). We therefore dubbed this the core region and measured the relative C_sat_ to be fourfold lower than entire NTR. The relative activity of the other polypeptides, as measured by C_sat,_ correlated with the fraction of core sequence present in each truncation. Despite the apparent sufficiency of the core, no single sequence stretch within the core is absolutely necessary, as illustrated by truncations NTR_100-250_ and NTR_250-400_. Both are capable of forming RNA-dependent condensates but contain non-overlapping segments of the core. Intriguingly, a fragment that lacked the N-terminal sequences but retained the core and acidic C-terminal sequences also appeared incapable of forming RNA-dependent condensates (Fig. 3A, B).

**Figure 3.**
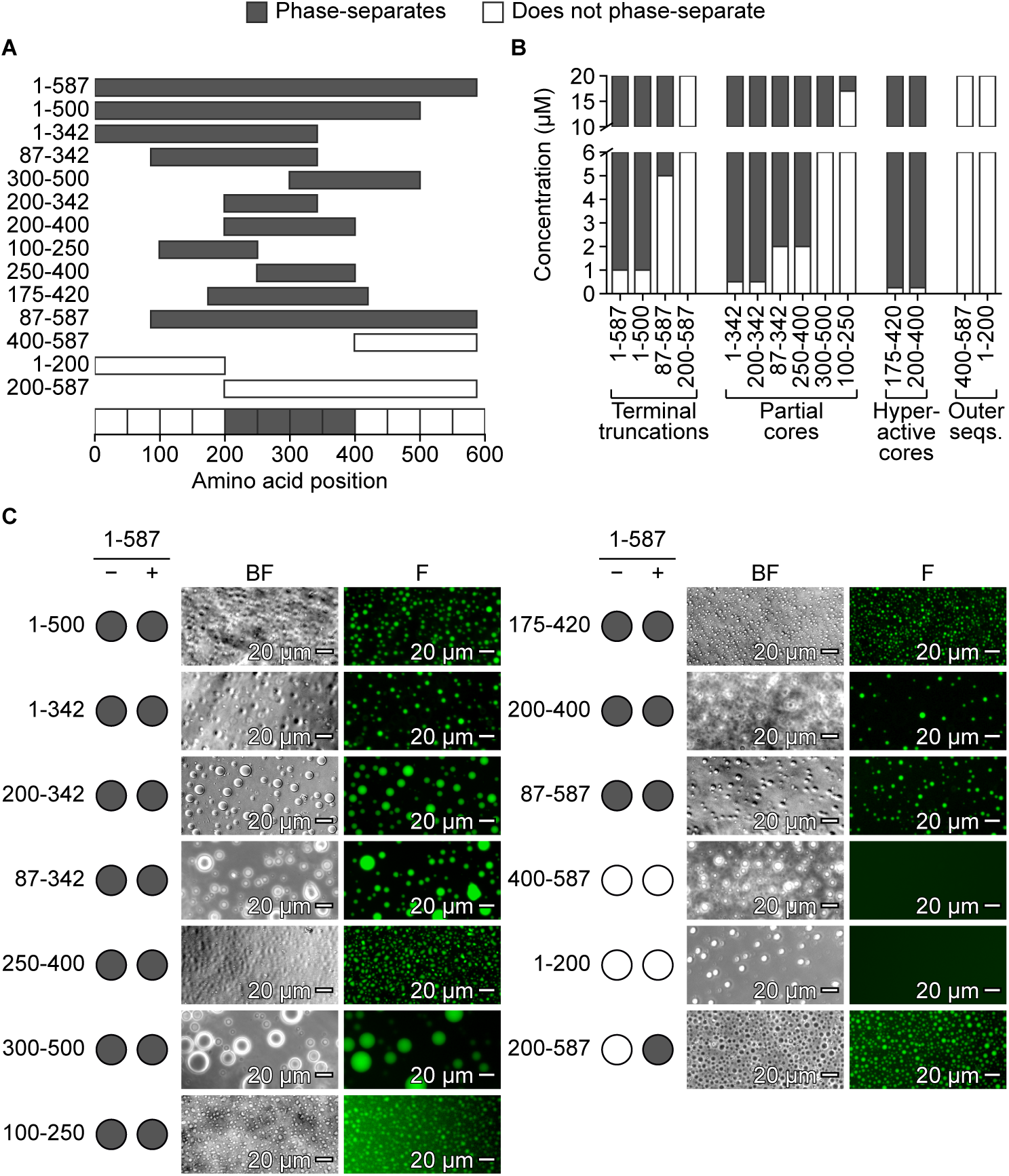
A central core of the eIF5B-NTR drives phase separation. **(A)** Schematic of the eIF5B-NTR truncations tested for RNA-dependent phase separation, with residue boundaries on the left. Grey bars indicate phase-separation and white bars indicate no phase-separation. The bottom bar summarizes phase-separation activity across the NTR in 50-residue windows and identifies residues 200-400 as the core. **(B)** Saturation concentrations (C_sat_) of eIF5B-NTR truncations. Each bar spans the protein concentrations tested up to 20 µM. White marks concentrations with no droplets and grey marks concentrations at which droplets formed; the grey-white boundary gives the C_sat_. **(C)** Partitioning assay in which each mNG-tagged truncation (66 nM, 2%) was added to condensates formed by full-length eIF5B-NTR (3.3 µM) and RNA (20 ng/µl). Circles indicate whether the truncation formed condensates on its own (previous titration to 20 µM) (−) or partitioned into full-length condensates (+); grey, yes; white, no. Representative brightfield (BF) and fluorescence (F) images of the + condition are shown. Results were scored as positive if droplets were fluorescent in at least two of three independent replicates. Scale bars, 50 µm.

To test the origin of the lack of activity of some of our truncations, we developed a partitioning assay. The purpose of this assay is to assess whether inactive truncations were still capable of forming the protein-protein interactions necessary to enter the droplet. In our binary system, the ability of a protein to form condensates requires the truncation to satisfy a sufficiently high valency of protein-protein interactions as well as protein-RNA interactions. To decipher which type of interaction was deficient in our inactive truncations, we formed condensates using the unlabeled full-length eIF5B-NTR and RNA as described above. To these droplets, we added a low concentration (2% mol/mol) of an mNG-tagged version of each truncation. If this truncation can form protein-protein interactions with the full-length NTR, it should be enriched into the full-length eIF5B-NTR droplets. Unsurprisingly, all polypeptides capable of forming RNA-driven condensates maintained sufficient protein-protein interactions to form fluorescent condensates (Fig. 3C). In contrast, the peripheral sequences (1-200 and 400-587), which failed to form RNA-driven condensates, also failed to partition with full-length NTR. Suggesting that these sequences simply cannot form the required protein-protein interactions. By contrast, the N-terminal truncation NTR_200-587,_ which failed to form RNA-driven condensates, partitioned into the full-length condensates. Indicating that this polypeptide can form the interactions requisite for entering the phase but cannot form the phase on its own, at least at the concentrations tested here.

### Flanking regions modulate condensate material properties

Truncation profiling of the eIF5B-NTR identified a minimal core that is sufficient to initiate phase separation. Since a substantial fraction of the eIF5B-NTR is dispensable for this activity, we asked whether the flanking sequences instead are capable of tuning the material properties of the resulting condensates. To evaluate this, we used our tiled NTR polypeptides to study the properties of the full-length NTR versus the core and how adding peripheral sequences would modulate these properties. The fluidity of the core, as measured by FRAP, is drastically more static with a half-recovery time of 60.5 s compared to 1.3 s for the entire NTR (Fig. 4A). This led us to hypothesize that the extremities of the disordered protein may function to make the resulting condensate more fluid. Intermediate length truncations support this trend with recovery times between that of the stationary core and the highly dynamic entire NTR (Fig. 4A).

**Figure 4.**
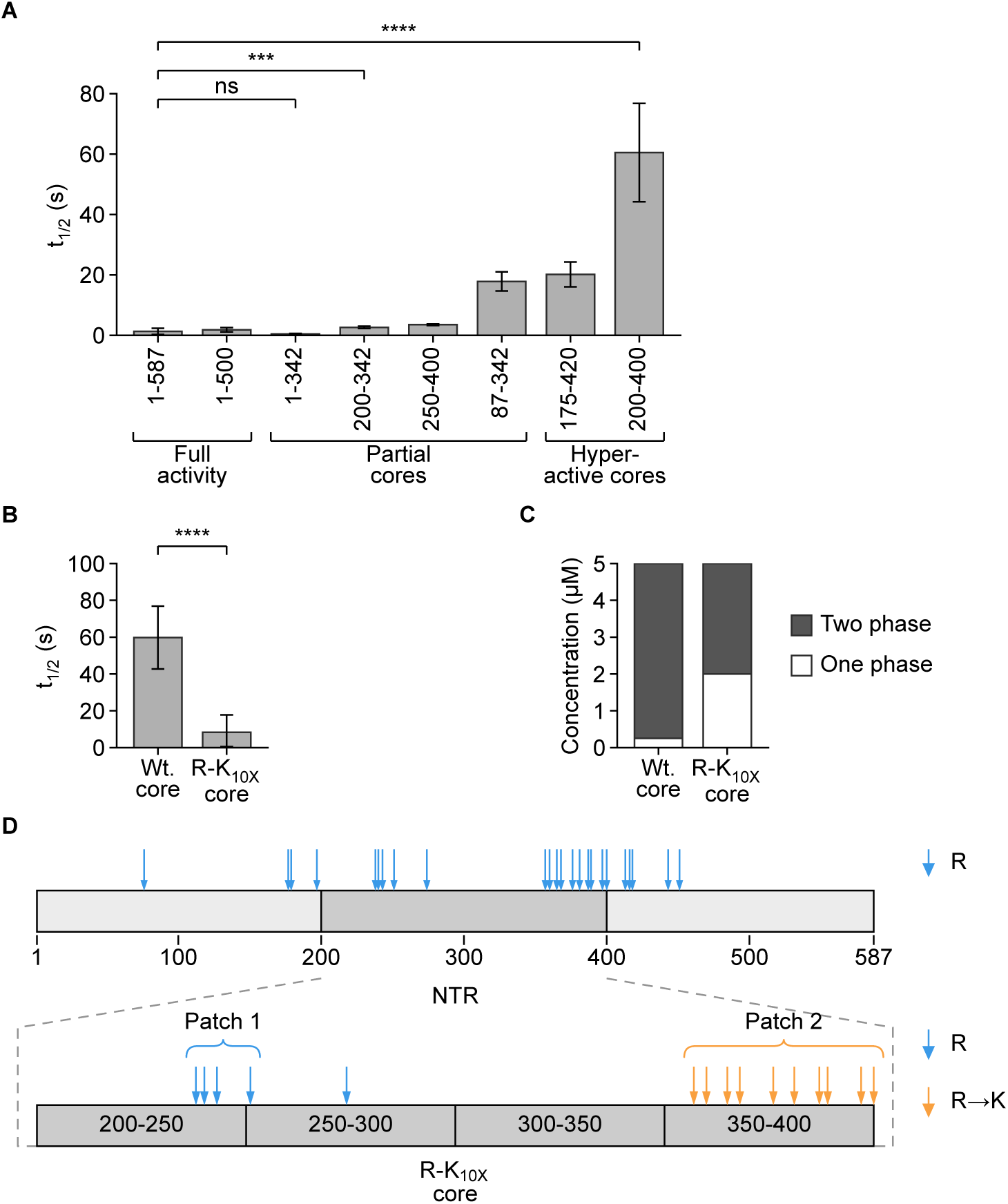
Arginine enrichment in the core controls condensate rigidity and phase-separation propensity. **(A)** FRAP recovery times (t_1/2_) of condensates formed by eIF5B-NTR truncations, grouped as full activity (1-587, 1-500), partial cores (1-342, 200-342, 87-342, 250-400) and hyperactive cores (200-400, 175-420). **(B)** Recovery time of condensates formed by the wild-type core (NTR_200-400_) and the R-K_10x_ core, in which ten arginine residues are mutated to lysine. **(C)** C_sat_ of the wild-type and R-K_10x_ cores. Grey marks protein concentrations giving two phases and white marks concentrations giving one phase. **(D)** Schematic of the NTR core (residues 200-400) showing arginine positions. Arginine in patch 1 and elsewhere in the core are shown in blue; those in patch 2 that were substituted with lysine in the R-K_10x_ core are shown in orange (R→K). Data in A and B are mean ± SD of 15 droplets from three independent replicates. Unpaired t test: ns, p ≥ 0.05; ***, p < 0.001; ****, p < 0.0001.

Previous work from multiple labs has demonstrated that increasing the ratio of arginine to lysine in a phase-separating peptide significantly increases the resulting viscosity of the condensate as well as stabilizes the RNA-protein complex (42, 43). Therefore, a plausible basis for the relative rigidity of the core is its enrichment in arginine residues. Arginines are concentrated in two patches spanning R238-R251 (5 arginine) and R357-R418 (13 arginine). Notably these patches overlap well with the eIF5B-NTR core (Fig. 4D). While the entire NTR is a low complexity polyampholyte, lysine predominates as the primary cationic amino acid with 83% of these charged residues being lysine relative to 16% arginine. In contrast, in the core region, arginine accounts for 22% of cationic residues.

To assess whether the arginine-to-lysine ratio was substantively regulating condensate properties, we generated a variant of the core in which ten arginine residues in the second arginine patch had been changed to lysine (200-400R-K_10X_). This effectively maintains the net charge properties while lowering the arginine-to-lysine ratio in the core. When fluidity was assessed using FRAP, the t_1/2_ of the R-K_10X_ core was 9.1 s (Fig. 4B). Relative to the wild-type core t_1/2_ of 60.5 s, this mutant was substantially more fluid, consistent with the arginine-to-lysine ratio acting as a determinant of condensate fluidity. Notably, the same loss of arginine residues also greatly increased the C_sat_ of the core from 0.25 µM to 2 µM, demonstrating a reduction in the total phase-separation propensity of the polypeptide (Fig. 4C).

### Cellular studies to support the in vitro model

Our minimal in vitro system establishes that eIF5B-NTR is sufficient to form fluid, RNA-dependent condensates in near physiological conditions. But our reconstituted binary system does not recapitulate the crowded and heterogeneous cellular milieu, so we next examined the behavior of eIF5B-NTR condensates in cells. Previously, we showed that overexpression of full length eIF5B-NTR is sufficient to drive condensate formation in cells (28). To characterize these structures, we overexpressed an mNG-eIF5B-NTR fusion and performed FRAP on the resulting granules (Fig. 5A-C). The rapid recovery (t_1/2_ = .15 s) indicates a highly mobile, liquid-like body, consistent with the eIF5B-NTR forming dynamic liquid-liquid phase condensates in a cellular environment.

**Figure 5.**
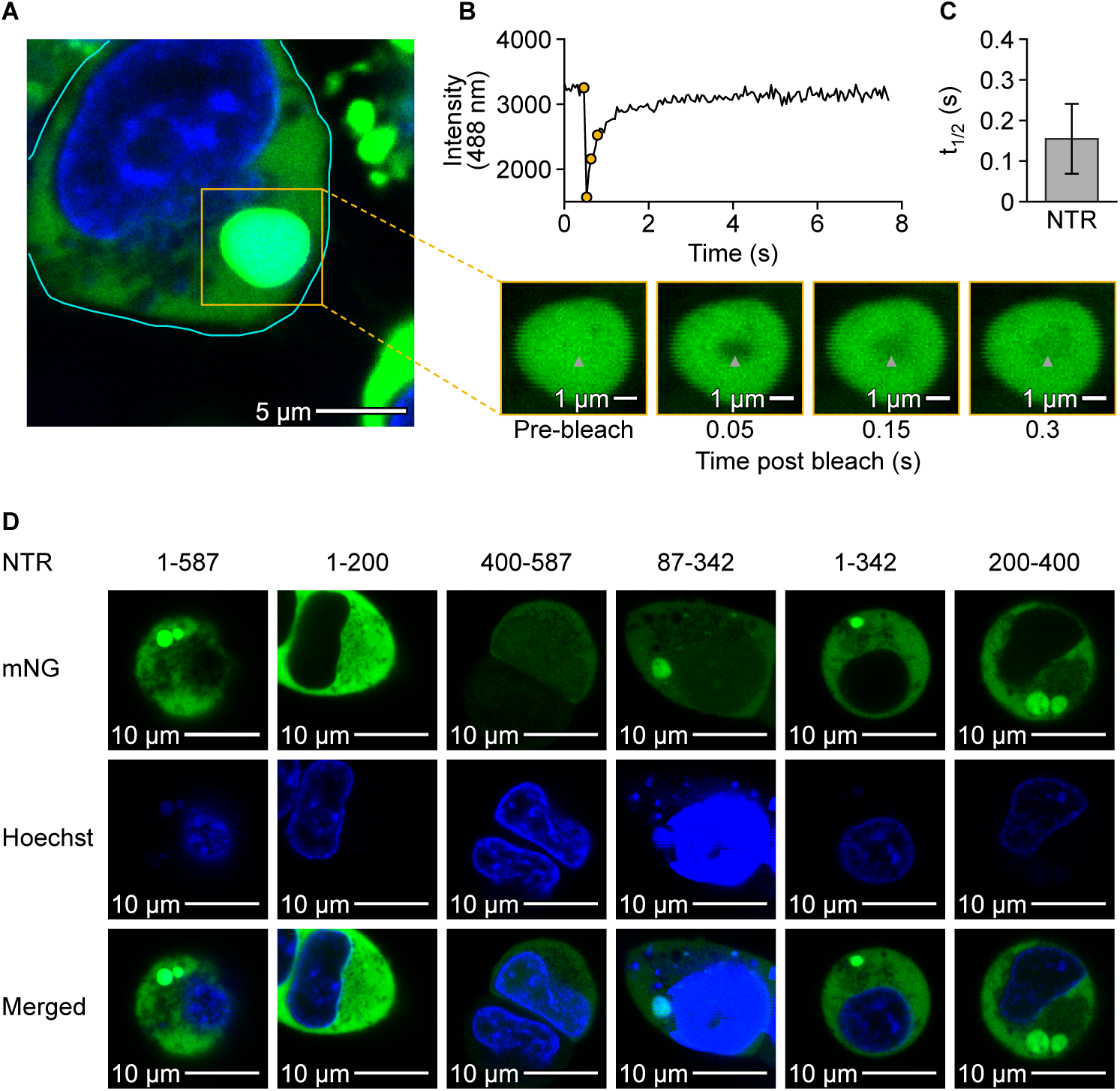
eIF5B-NTR forms dynamic, core-dependent condensates in cells. **(A)** Confocal image of a HEK293T cell expressing mNG-eIF5B-NTR (green), counterstained with Hoechst (blue), 48 hr. after transfection. The cyan line outlines the cell. Scale bar, 5 µm. **(B)** Representative FRAP recovery trace of a cytoplasmic mNG-eIF5B-NTR granule. Images below the trace show the granule before and 0.05, 0.15 and 0.3 s after the bleach; the arrowhead marks the bleached region. Scale bars, 1 µm. **(C)** Recovery times (t½) of mNG-eIF5B-NTR granules in cells. The bar shows mean ± SD (n = 23 granules from three independent transfection). **(D)** Representative confocal images of HEK293T cells expressing mNG-tagged eIF5B-NTR truncations 48 hr. after transfection, with rows for mNG, Hoechst and the merged image. All images are shown at the same scale, without intensity adjustment. Scale bars, 10 µm.

To further test our model of eIF5B-NTR as a phase separation scaffold, we decided to validate the ability of key eIF5B-NTR truncations to form condensates in a cellular system. To do so, we cloned truncated versions of mNG-fused eIF5B-NTR under the constitutive expression of a CMV promoter and transiently expressed each construct in HEK293T cells. When necessary, a second fluorescent tag was added to ensure nuclear exclusion and prevent formation of nuclear granules (Fig. S3A). Formation of cellular granules was assayed 48 hours post-transfection by confocal microscopy (Fig. 5D). Cellular expression levels were assayed by measuring total fluorescence of cell lysates (Fig. S3B). This was done to ensure any peptide that failed to form granules was still expressing comparable levels of protein. Total fluorescence of NTR_1-200_ and NTR_400-587_ were both within a 2.5-fold range of full-length NTR expression and well above the poorest expressor, NTR_1-342_ which continued to form granules. Our imaging results are fully consistent with our in vitro model. Truncations that contain the eIF5B-NTR core (200–400) are capable of forming cellular condensates upon overexpression, while peripheral sequences failed to form condensates.

## Discussion

Our results demonstrate that the intrinsically disordered N-terminal region of eIF5B (NTR) is capable of liquid-liquid phase separation and that the NTR undergoes RNA-dependent phase separation under physiologically relevant conditions. This behavior is primarily driven by electrostatic interactions and can be modulated by both RNA concentration and RNA structure. This offers a potentially intriguing mechanism by which eIF5B may undergo condensation in the event of cellular stress. Namely, during stress events, the increased concentration of non-translating messenger RNA can outcompete the cis-electrostatic interactions within the intrinsically disordered domain of eIF5B, thereby forming multivalent networks. This potential mechanism parallels the proposed trigger and activity of G3BP1 in forming stress granules (44). Intriguingly, generic polyanions are also sufficient to induce condensation supporting the conclusion that the condensate is driven by the electrostatic interactions with the RNA and not nucleic acid sequence.

Against the backdrop of an increasingly well-defined molecular grammar for biomolecular condensation, the NTR of eIF5B is an instructive example of a naturally occurring, intrinsically disordered, polyampholyte (Fig. 6). Remarkably, despite the length and high charge density of the full NTR, only a compact segment is both necessary and sufficient for condensation, and the enrichment of arginine at this activity center points to a coherent mechanistic model. In this view, the two arginine patches of the core supply the high-valency contacts that drive robust condensation but yield rigid assemblies with slow dynamics. The flanking regions counterbalance this rigidity: their lysine-rich composition lowers the arginine-to-lysine ratio and restores fluidity, at the cost of condensation strength (Table S1). The partial and complete loss of activity in N-terminal truncations, NTR_87-587_ and NTR_200-587_ suggest that the N-terminal flank is important in providing charge or valency needed to form condensates when the highly acidic C-terminal stretch of the NTR is present. The loss of N-terminal sequence only appears vital in context of the complete acidic C-terminal sequence as can be seen by the robust activity of NTR_200-342_ and NTR_200-400_. Taken together, the material state of the condensate thus emerges from a tuned competition of sequence identity which appears to have been patterned into functional regions.

**Figure 6.**
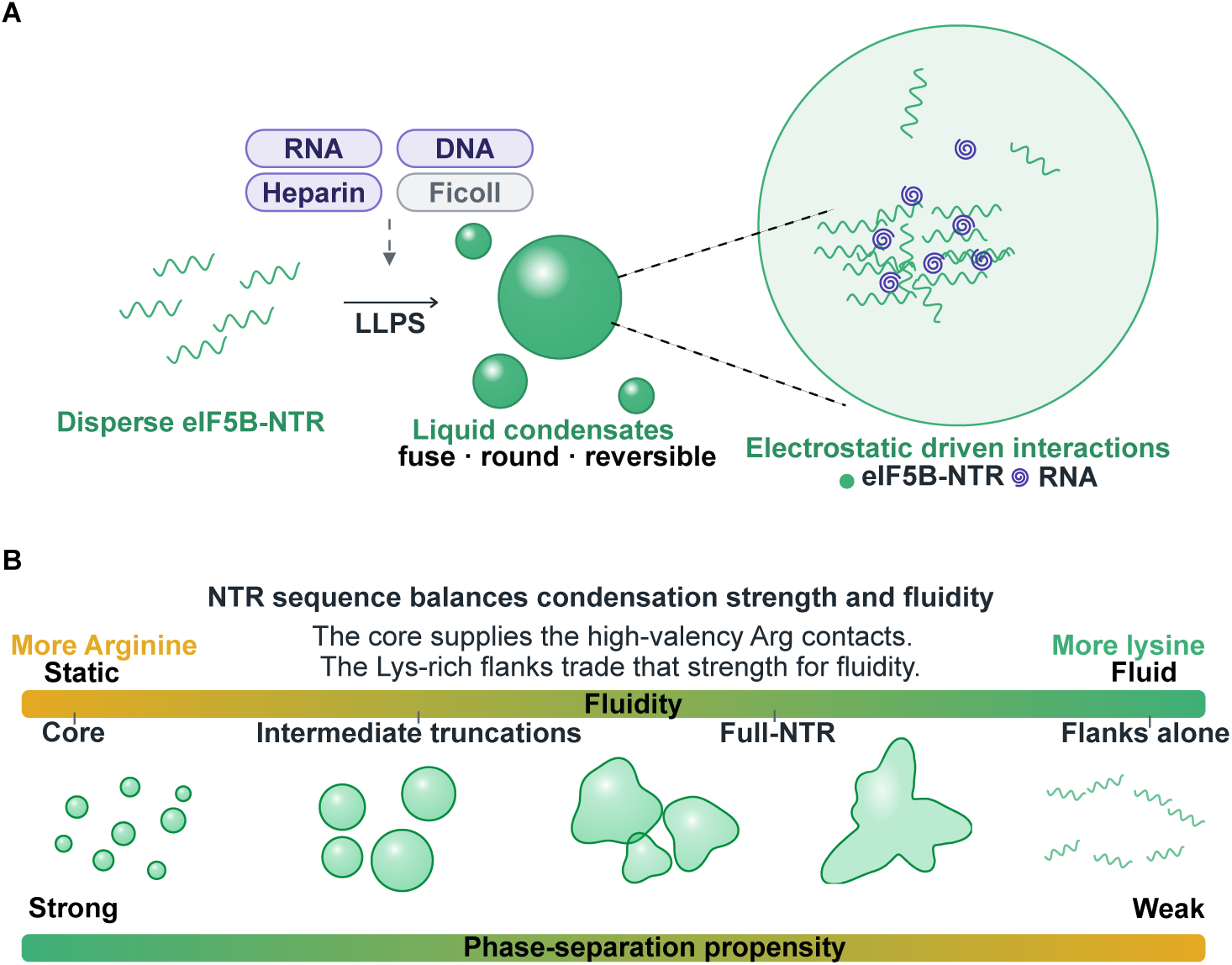
Model of eIF5B-NTR phase separation. **(A)** Cartoon depiction of eIF5B-NTR’s interaction with condensate partners to form multivalent interactions. Subset depicts the network of RNA-protein interactions that are hypothesized to drive condensation in vivo. **(B)** Depicts the sequence trade-off implied by core versus flanking sequences of the NTR. Sequence containing the Arginine rich core form strong but static condensates while flanking regions alone form interactions that are too weak to condensate. NTR depicted as balancing these two extremes.

The core-flanks architecture conveys regulatory logic. Flanks tempering the intrinsic phase-separation propensity of eIF5B may be essential to prevent premature, stress-independent conversion from a dispersed state into granules. Because the cell contains a concentration of eIF5B close to the saturation concentration, the conversion to granules can be easily triggered through increased free RNA in the advent of cellular stress (45). Alternatively, the dysregulation of eIF5B concentrations, such as in cancer, may provide an alternative signal for granule formation (33). This would provide a mechanistic link between the non-canonical translation that drives poor patient prognosis to the dysregulation of granule formation. A further layer of control is suggested by the extensive, dynamic phosphorylation that decorates this disordered region and could rapidly tune the charge balance that governs both condensation and fluidity (46–48). Together, these observations converge on a model in which a minimal, arginine-driven core provides the latent capacity to condense, while the flanking lysine-rich sequences of the disordered N-terminus act as tunable rheostats of physiological cues such as stress and phosphorylation. Defining how these elements are integrated in vivo will be an important next step toward understanding how this translation factor is repurposed as a condensate scaffold under conditions of cellular stress and disease.

## Experimental procedures

### Sequence property and diagramming

Sequence analysis generated using the MolPhase from the MIAO Yansong Lab, from Nanyang Technological University (https://molphase.sbs.ntu.edu.sg/).

### Cloning and construct preparation

All newly generated plasmid constructs were made using New England Biolabs’ HiFi DNA assembly kit as described by the manufacturer. The expression plasmid is based on the pET28 vector (Novagen/Merck). Due to a high degree of truncated peptide products during expression of this protein, we found that using a 6x-his tag on the C-terminus of all bacterial constructs provided the easiest method for expression and purification. All constructs were fully sequenced using whole-plasmid sequencing before use.

### Bacterial expression and purification

To purify all proteins, we co-transformed the relevant plasmid with the pRARE2 plasmid containing rare codon tRNAs into a BL21 Star (DE3) bacterial strain. Single colonies were selected from a plate with the applicable antibiotic and grown overnight at 37 °C. Expression cultures were made by diluting the overnight culture 1 to 100 and allowed to grow to an OD_600_ of 0.5 before induction with 1 mM IPTG. Post-induction cultures were shifted from 37 °C to 28 °C and allowed to express overnight.

### Protein purification

Purification was accomplished utilizing a three-step purification scheme. Briefly, bacterial pellets were resuspended in a high salt nickel purification buffer A (25 mM Tris pH 7.5, 750 mM NaCl, 25 mM imidazole, 1 mM EDTA, 2 mM DTT). High salt was maintained through the initial nickel purification to prevent premature phase separation. Samples were lysed using a microfluidizer and clarified via centrifugation. The soluble fraction was applied to Ni Sepharose 6 Fast Flow via batch loading and washed with nickel buffer A. Proteins were eluted with a nickel elution buffer (25 mM Tris, pH 7.5, 750 mM NaCl, 500 mM imidazole, 1 mM EDTA, 2 mM DTT). Protein concentration was checked with a Bradford assay prior to desalting into a low salt, heparin-binding buffer (25 mM Tris pH 7.5, 100 mM NaCl, 2 mM DTT). This solution was applied to a heparin Sepharose 6 FF column and eluted using a linear 20 column volume gradient with heparin buffer B (25 mM Tris, pH 7.5, 1M NaCl, 2 mM DTT). Fractions were characterized via SDS-PAGE; fractions containing the correct product were polished on a Sephacryl S-300 HR, 120ml size exclusion column. All proteins were stored with the final addition of 50% glycerol and stored at −20 °C. Working concentrations were determined by quantitative SDS-PAGE after desalting into assay buffer.

### In vitro transcription (IVT) of RNA

RNAs were synthesized by generating linear DNA templates via PCR with the addition of a T3 phage promoter. 100 µl IVT reactions were assembled with (40 mM Tris-HCL, pH 7.9, 10 mM DTT, 2 mM spermidine-HCL, 20 mM Mg(OAc)_2_, 6 mM rNTPs, 0.1 µg/µl template and 500 units T3 RNAP). Reactions were run for 30 min at 37°C prior to quenching and precipitation using EDTA/lithium chloride. Collected RNA pellets were washed with 80% ethanol twice before resuspension in ddH_2_O. RNAs were checked via TBE-urea agarose with ethidium bromide staining for purity and DNA contamination before use in assays.

**Table 1.**
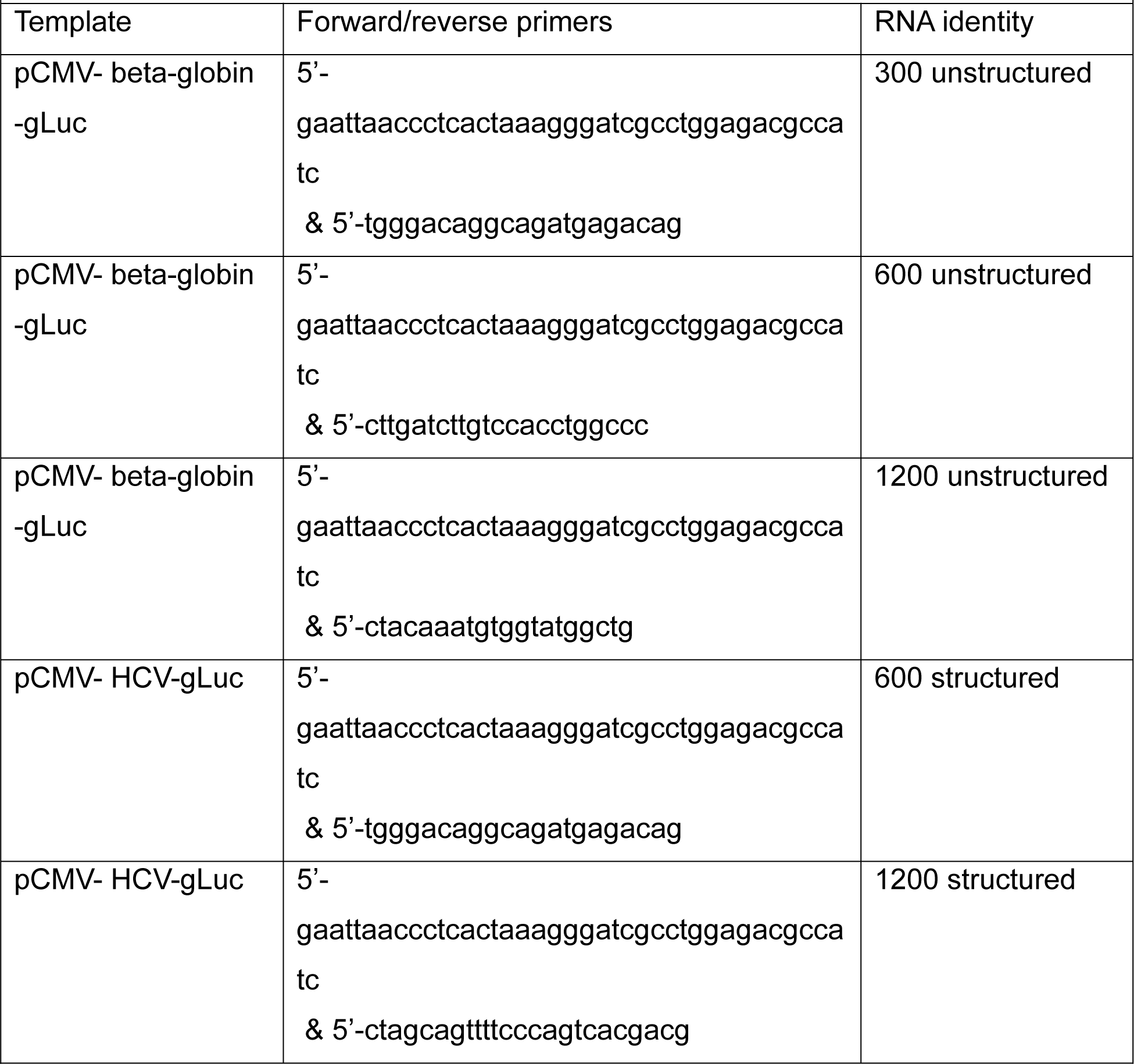
Primers for making IVT templates.

### Fluorescent RNA labeling for two-color FRAP

RNA was fluorescently labeled with a hydrazide conjugated 594CF™ dye at its 3′ end. To carry out this reaction, RNA was first diluted into 100 mM sodium acetate, pH 4.5. The terminal ribose was oxidized by the addition of 10 mM sodium periodate with rotation at 4 °C for 1 hr. Oxidized RNA was precipitated with 300 mM sodium acetate, pH 5.2, and 3 volumes of ethanol. Dried and washed RNA pellets were resuspended in ddH_2_O and conjugated with a hydrazide-functionalized fluorophore. This reaction was allowed to proceed overnight, after which unincorporated dye was removed and labeled RNA was recovered by ethanol precipitation. An additional GE-microspin desalting column was added to aid in the complete removal of free dye. Finally, labeling was confirmed by comparing labeled and unlabeled RNA by TBE-agarose gel electrophoresis prior to use.

### Phase separation assay

Phase separation was monitored by an imaging-based assay. The test protein was desalted from storage buffer into assay buffer (25 mM Tris pH 7.5, 100 mM NaCl, 2 mM MgCl_2_, 1 mM DTT, unless otherwise specified). A separately desalted mNG-tagged tracer of the same protein was added at 2% of the untagged protein, followed by IVT-synthesized RNA (300 nucleotides unless otherwise noted) to a final concentration of 20 ng/µl. Reactions were equilibrated at room temperature for 30 min, after which an aliquot was transferred to an imaging well and centrifuged briefly (400 RCF, 1 min) to settle droplets onto the imaging surface.

### Ficoll Titrations

To identify the conditions under which eIF5B-NTR undergoes Ficoll dependent phase separation, we performed a two-dimensional titration varying both the molecular crowding agent (Ficoll) and the protein concentration. Ficoll PM 400 was prepared as a concentrated stock (e.g., 40% w/v) in assay buffer 25 mM Tris pH 7.5, 100 mM NaCl, 2 mM MgCl_2_, 1 mM DTT. Protein was buffer exchanged into the same assay buffer and 2% mNG-eIF5B-NTR was added for visualization. All components were combined at room temperature with Ficoll titrated across a range of final concentrations (e.g., 0, 2.5, 5, 10, 15, and 20% w/v) against a series of protein concentrations from 0.25 µM to 3 µM. Each condition was assembled in a PCR tube prior to transferring to a low volume 96-well plate and imaging at 20x magnification on a Keyence BZX710 fluorescent microscope. All conditions were performed in three independent replicates, and representative images were acquired 30 min after mixing.

### Protein-RNA titrations and C_sat_ measurements

To measure the permissive conditions for phase separation protein and RNA were varied into 2-fold dilutions prior to RNA droplet nucleation. Only after dilution were protein and RNA combined and tested with the phase separation assay as described above. The C_sat_ of a protein was defined as the lowest concentration of protein in which at least 10 droplets were clearly visible. All titration experiments were conducted in triplicate and imaged in a reduced volume 96 well-plate with a Keyence BZX710 fluorescent microscope.

### Small molecule testing

To test the ability of small molecules to inhibit RNA-dependent phase separation we performed phase separation assays as described above. To each reaction the small molecule of interest was diluted to 2x concentration in buffer prior to the addition of protein. Protein and buffer containing small molecules were mixed well before the addition of RNA to nucleate droplet formation. All samples were performed in triplicate and imaged as described above.

### Partitioning assay

Partitioning assays were performed by generating samples containing 3.3 µM eIF5B-NTR and 20ng/µl IVT synthesized RNA in phase separation buffer 25 mM Tris pH 7.5, 100 mM NaCl, 2 mM MgCl_2_, 1 mM DTT. To this sample 66nM (2%) of the relevant mNG-eIF5B-NTR peptide was added to the solution and allowed to come to equilibrium prior to the solution being added to reduced volume 96 well plate and imaged on a Keyence BZX710 fluorescent microscope with multi-channel settings to capture both brightfield and fluorescent images. All peptides were tested in triplicate and determined to partition if droplets were clearly distinguished in the fluorescent channel in at least 2 of 3 replicates.

### Droplet FRAP and analysis

FRAP measurements were taken by generating droplets as previously described. Droplet containing solutions were then applied to imaging slides pretreated with 1% Pluronic^®^ F-127 for 1 hour and then washed with 20% ethanol before air drying. Samples were contained on the slide using a Grace Bio-Labs silicone gasket with a coverslip added to prevent evaporation during imaging. Droplets were imaged and bleached using a Zeiss-LSM-880 confocal microscope utilizing the 488nm argon laser. All imaging samples were acquired using identical imaging parameters in triplicate with at least 5 droplets imaged per sample. Analysis of fluorescent recovery was digitized using ImageJ and normalized to droplet bleaching. Recovery curves were fitted to a simple exponential using the curve fitting function in GraphPad Prism.

### Cell culture and transfections

HEK293T cells (human embryonic kidney 293 cells stably expressing the SV40 large T antigen; ATCC CRL-3216, RRID:CVCL_0063) were obtained from ATCC. All cell cultures were maintained using aseptic practices in 10% FBS supplemented DMEM - high glucose with sodium bicarbonate, ʟ-glutamine, sodium pyruvate and a 1x Penicillin-Streptomycin cocktail. Cell lines were maintained under 20 passages and transiently transfected using Polyethyleneimine. Cells were transfected between 20-50% confluency and media was changed 12 hours post transfection. Cells were then allowed to grow for 48 hours post transfection prior to imaging.

### Cellular FRAP

For cellular FRAP, cells were transiently transfected as described and counterstained with Hoechst 33342 30 min before imaging. Imaging and photobleaching were performed on a Zeiss LSM 880 confocal microscope using the 488 nm argon laser. Experiments were performed in triplicate, with 5 cells bleached and measured per replicate. Recovery curves were digitized in ImageJ, fitted to a single exponential function, and t_1/2_ values were calculated in GraphPad Prism.

### Granule formation and expression profiling

Cytoplasmic granule formation was assessed following transient transfection as described above. Transfected cells were either transferred to a sterile 96-well plate for expression profiling or to a chambered coverslip for imaging. For expression profiling, cells were washed twice with PBS 48 hours after transfection and lysed directly in the plate on ice for 10 min (50 mM Tris pH 7.5, 150 mM NaCl, 1% IGEPAL). Total fluorescence of lysates was read on a Tecan plate reader with GFP filters, with eight parallel replicates per sample. For granule imaging, cells on coverslips were stained with 0.5x Hoechst 33342 30 min before acquisition on a Zeiss LSM 880 confocal microscope; each field was first captured as a single-plane image for droplet identification and then as a z-stack spanning the full-depth of the cell. All samples were imaged in three independent replicates with at least 3 representative cells per sample per replicate.

### Statistical analysis

Unless otherwise noted, all experiments were performed with at least three independent replicates. Data are reported as mean ± SD. Statistical comparisons were performed in GraphPad Prism using unpaired-t tests, with ns; p ≥0.05, *; p< 0.05, ** p<0.01, ***; p<0.001, ****; p<0.0001

## Data availability

All data are contained within the manuscript and its supporting information.

## Supporting information

This article contains supporting information.

## Acknowledgments.

“We acknowledge the use of the Brandeis University Light Microscopy Core Facility (RRID:SCR_025892) for instrumentation and support.

## Abbreviations

C_sat_, saturation concentration; CTD, C-terminal domain; eIF5B, eukaryotic initiation factor 5B; FCR, fraction of charged residues; FRAP, fluorescence recovery after photobleaching; HCV, hepatitis C virus; IDR, intrinsically disordered region; LCR, low-complexity region; MLO, membraneless organelle; NCPR, net charge per residue; NTR, N-terminal region; t_1/2,_ half-time of recovery.

## Reference

1. Banani, S. F., Lee, H. O., Hyman, A. A., and Rosen, M. K. (2017) Biomolecular condensates: organizers of cellular biochemistry Nat Rev Mol Cell Biol 18, 285–298

2. Hirose, T., Ninomiya, K., Nakagawa, S., and Yamazaki, T. (2023) A guide to membraneless organelles and their various roles in gene regulation Nature Reviews Molecular Cell Biology 24, 288–304

3. Shin, Y., and Brangwynne, C. P. (2017) Liquid phase condensation in cell physiology and disease Science 357, eaaf4382

4. Ausserwoger, H., de Csillery, E., Qian, D., Krainer, G., Welsh, T. J., Sneideris, T., et al. (2025) Quantifying collective interactions in biomolecular phase separation Nat Commun 16, 7724

5. Sun, J., Qu, J., Zhao, C., Zhang, X., Liu, X., Wang, J., et al. (2024) Precise prediction of phase-separation key residues by machine learning Nature Communications 15,

6. Dignon, G. L., Best, R. B., and Mittal, J. (2020) Biomolecular Phase Separation: From Molecular Driving Forces to Macroscopic Properties Annu Rev Phys Chem 71, 53–75

7. Elbaum-Garfinkle, S., and Kriwacki, R. (2025) Biomolecular Condensates; Emerging Themes and Concepts J Mol Biol 437, 169449

8. Pak, C. W., Kosno, M., Holehouse, A. S., Padrick, S. B., Mittal, A., Ali, R., et al. (2016) Sequence Determinants of Intracellular Phase Separation by Complex Coacervation of a Disordered Protein Mol Cell 63, 72–85

9. Poh, C. W., and Mueller-Cajar, O. (2024) π-π Interactions Drive the Homotypic Phase Separation of the Prion-like Diatom Pyrenoid Scaffold PYCO1 J Mol Biol 436, 168800

10. Qamar, S., Wang, G., Randle, S. J., Ruggeri, F. S., Varela, J. A., Lin, J. Q., et al. (2018) FUS Phase Separation Is Modulated by a Molecular Chaperone and Methylation of Arginine Cation-π Interactions Cell 173, 720–734 e715

11. Nott, T. J., Petsalaki, E., Farber, P., Jervis, D., Fussner, E., Plochowietz, A., et al. (2015) Phase transition of a disordered nuage protein generates environmentally responsive membraneless organelles Mol Cell 57, 936–947

12. Banjade, S., Wu, Q., Mittal, A., Peeples, W. B., Pappu, R. V., and Rosen, M. K. (2015) Conserved interdomain linker promotes phase separation of the multivalent adaptor protein Nck Proc Natl Acad Sci U S A 112, E6426–6435

13. Li, P., Banjade, S., Cheng, H.-C., Kim, S., Chen, B., Guo, L., et al. (2012) Phase transitions in the assembly of multivalent signalling proteins Nature 483, 336–340

14. Elbaum-Garfinkle, S., Kim, Y., Szczepaniak, K., Chen, C. C., Eckmann, C. R., Myong, S., et al. (2015) The disordered P granule protein LAF-1 drives phase separation into droplets with tunable viscosity and dynamics Proc Natl Acad Sci U S A 112, 7189–7194

15. Yang, P., Mathieu, C., Kolaitis, R.-M., Zhang, P., Messing, J., Yurtsever, U., et al. (2020) G3BP1 Is a Tunable Switch that Triggers Phase Separation to Assemble Stress Granules Cell 181, 325–345.e328

16. Owen, I., and Shewmaker, F. (2019) The Role of Post-Translational Modifications in the Phase Transitions of Intrinsically Disordered Proteins Int J Mol Sci 20,

17. Pederson, T. (2011) The nucleolus Cold Spring Harb Perspect Biol 3,

18. Brown, D. D., and Gurdon, J. B. (1964) ABSENCE OF RIBOSOMAL RNA SYNTHESIS IN THE ANUCLEOLATE MUTANT OF XENOPUS LAEVIS Proc Natl Acad Sci U S A 51, 139–146

19. Kass, S., Tyc, K., Steitz, J. A., and Sollner-Webb, B. (1990) The U3 small nucleolar ribonucleoprotein functions in the first step of preribosomal RNA processing Cell 60, 897–908

20. Koberna, K., Malínský, J., Pliss, A., Masata, M., Vecerova, J., Fialová, M., et al. (2002) Ribosomal genes in focus: new transcripts label the dense fibrillar components and form clusters indicative of “Christmas trees” in situ J Cell Biol 157, 743–748

21. Brangwynne, C. P., Mitchison, T. J., and Hyman, A. A. (2011) Active liquid-like behavior of nucleoli determines their size and shape in Xenopus laevis oocytes Proceedings of the National Academy of Sciences 108, 4334–4339

22. Tourrière, H., Chebli, K., Zekri, L., Courselaud, B., Blanchard, J. M., Bertrand, E., et al. (2003) The RasGAP-associated endoribonuclease G3BP assembles stress granules J Cell Biol 160, 823–831

23. Kedersha, N. L., Gupta, M., Li, W., Miller, I., and Anderson, P. (1999) RNA-binding proteins TIA-1 and TIAR link the phosphorylation of eIF-2 alpha to the assembly of mammalian stress granules J Cell Biol 147, 1431–1442

24. Brangwynne, C. P., Eckmann, C. R., Courson, D. S., Rybarska, A., Hoege, C., Gharakhani, J., et al. (2009) Germline P Granules Are Liquid Droplets That Localize by Controlled Dissolution/Condensation Science 324, 1729–1732

25. Patel, A., Lee, H. O., Jawerth, L., Maharana, S., Jahnel, M., Hein, M. Y., et al. (2015) A Liquid-to-Solid Phase Transition of the ALS Protein FUS Accelerated by Disease Mutation Cell 162, 1066–1077

26. Molliex, A., Temirov, J., Lee, J., Coughlin, M., Kanagaraj, A. P., Kim, H. J., et al. (2015) Phase separation by low complexity domains promotes stress granule assembly and drives pathological fibrillization Cell 163, 123–133

27. Banani, S. F., Afeyan, L. K., Hawken, S. W., Henninger, J. E., Dall’Agnese, A., Clark, V. E., et al. (2022) Genetic variation associated with condensate dysregulation in disease Dev Cell 57, 1776–1788 e1778

28. Harris, M. T., and Marr, M. T., 2nd (2023) The intrinsically disordered region of eIF5B stimulates IRES usage and nucleates biological granule formation Cell Rep 42, 113283

29. Miao, E., Yang, D., Yue, X., Zhang, Z., Liu, H., Qin, H., et al. (2025) Revealing Stress Granule Compositional Heterogeneity through Antibody-Guided Proximity Labeling Anal Chem 97, 8313–8321

30. Youn, J.-Y., Dunham, W. H., Hong, S. J., Knight, J. D. R., Bashkurov, M., Chen, G. I., et al. (2018) High-Density Proximity Mapping Reveals the Subcellular Organization of mRNA-Associated Granules and Bodies Molecular Cell 69, 517–532.e511

31. Hubstenberger, A., Courel, M., Bénard, M., Souquere, S., Ernoult-Lange, M., Chouaib, R., et al. (2017) P-Body Purification Reveals the Condensation of Repressed mRNA Regulons Molecular Cell 68, 144–157.e145

32. Wang, Z. G., Zheng, H., Gao, W., Han, J., Cao, J. Z., Yang, Y., et al. (2016) eIF5B increases ASAP1 expression to promote HCC proliferation and invasion Oncotarget 7, 62327–62339

33. Suresh, S., Chen, B., Zhu, J., Golden, R. J., Lu, C., Evers, B. M., et al. (2020) eIF5B drives integrated stress response-dependent translation of PD-L1 in lung cancer Nat Cancer 1, 533–545

34. Ross, J. A., Dungen, K. V., Bressler, K. R., Fredriksen, M., Khandige Sharma, D., Balasingam, N., et al. (2019) Eukaryotic initiation factor 5B (eIF5B) provides a critical cell survival switch to glioblastoma cells via regulation of apoptosis Cell Death Dis 10, 57

35. Lapointe, C. P., Grosely, R., Sokabe, M., Alvarado, C., Wang, J., Montabana, E., et al. (2022) eIF5B and eIF1A reorient initiator tRNA to allow ribosomal subunit joining Nature 607, 185–190

36. Pestova, T. V., Lomakin, I. B., Lee, J. H., Choi, S. K., Dever, T. E., and Hellen, C. U. (2000) The joining of ribosomal subunits in eukaryotes requires eIF5B Nature 403, 332–335

37. Liang, Q., Peng, N., Xie, Y., Kumar, N., Gao, W., and Miao, Y. (2024) MolPhase, an advanced prediction algorithm for protein phase separation EMBO J 43, 1898–1918

38. Marianelli, A. M., Miller, B. M., and Keating, C. D. (2018) Impact of macromolecular crowding on RNA/spermine complex coacervation and oligonucleotide compartmentalization Soft Matter 14, 368–378

39. Bouchard, J. J., Otero, J. H., Scott, D. C., Szulc, E., Martin, E. W., Sabri, N., et al. (2018) Cancer Mutations of the Tumor Suppressor SPOP Disrupt the Formation of Active, Phase-Separated Compartments Mol Cell 72, 19–36 e18

40. Viegas, R. G., Sinha, A., Mainan, A., Sanbonmatsu, K. Y., Onuchic, J. N., Roy, S., et al. (2025) Critical Role of Mg(2+) Ions in RNA Folding Transitions: Anchoring the A-Minor Twist in the SAM-II Riboswitch J Phys Chem B 129, 9058–9067

41. Yamagami, R., Bingaman, J. L., Frankel, E. A., and Bevilacqua, P. C. (2018) Cellular conditions of weakly chelated magnesium ions strongly promote RNA stability and catalysis Nature Communications 9, 2149

42. Fisher, R. S., and Elbaum-Garfinkle, S. (2020) Tunable multiphase dynamics of arginine and lysine liquid condensates Nature Communications 11, 4628

43. Paloni, M., Bussi, G., and Barducci, A. (2021) Arginine multivalency stabilizes protein/RNA condensates Protein Sci 30, 1418–1426

44. Guillén-Boixet, J., Kopach, A., Holehouse, A. S., Wittmann, S., Jahnel, M., Schlüßler, R., et al. (2020) RNA-Induced Conformational Switching and Clustering of G3BP Drive Stress Granule Assembly by Condensation Cell 181, 346–361 e317

45. Hein, Marco Y., Hubner, Nina C., Poser, I., Cox, J., Nagaraj, N., Toyoda, Y., et al. (2015) A Human Interactome in Three Quantitative Dimensions Organized by Stoichiometries and Abundances Cell 163, 712–723

46. Olsen, J. V., Blagoev, B., Gnad, F., Macek, B., Kumar, C., Mortensen, P., et al. (2006) Global, in vivo, and site-specific phosphorylation dynamics in signaling networks Cell 127, 635–648

47. Dephoure, N., Zhou, C., Villen, J., Beausoleil, S. A., Bakalarski, C. E., Elledge, S. J., et al. (2008) A quantitative atlas of mitotic phosphorylation Proc Natl Acad Sci U S A 105, 10762–10767

48. Zhou, H., Di Palma, S., Preisinger, C., Peng, M., Polat, A. N., Heck, A. J. R., et al. (2012) Toward a Comprehensive Characterization of a Human Cancer Cell Phosphoproteome Journal of Proteome Research 12, 260–271

