## supplemental figures for "The intrinsically disordered N-terminal region of eIF5B nucleates RNA-driven phase separation"

### Contents

**Figure S1.** Purified eIF5B-NTR proteins and reversibility of eIF5B-NTR droplets. S-2

**Figure S2.** Tracer controls and condensate dynamics with 1200-nucleotide RNA. S-5

**Figure S3.** Cellular expression and granule formation of eIF5B-NTR truncations. S-6

**Table S1.** Physicochemical properties and phase-separation parameters of eIF5B-NTR constructs. S-7

**Figure S1**

**A** eIF5B-NTR

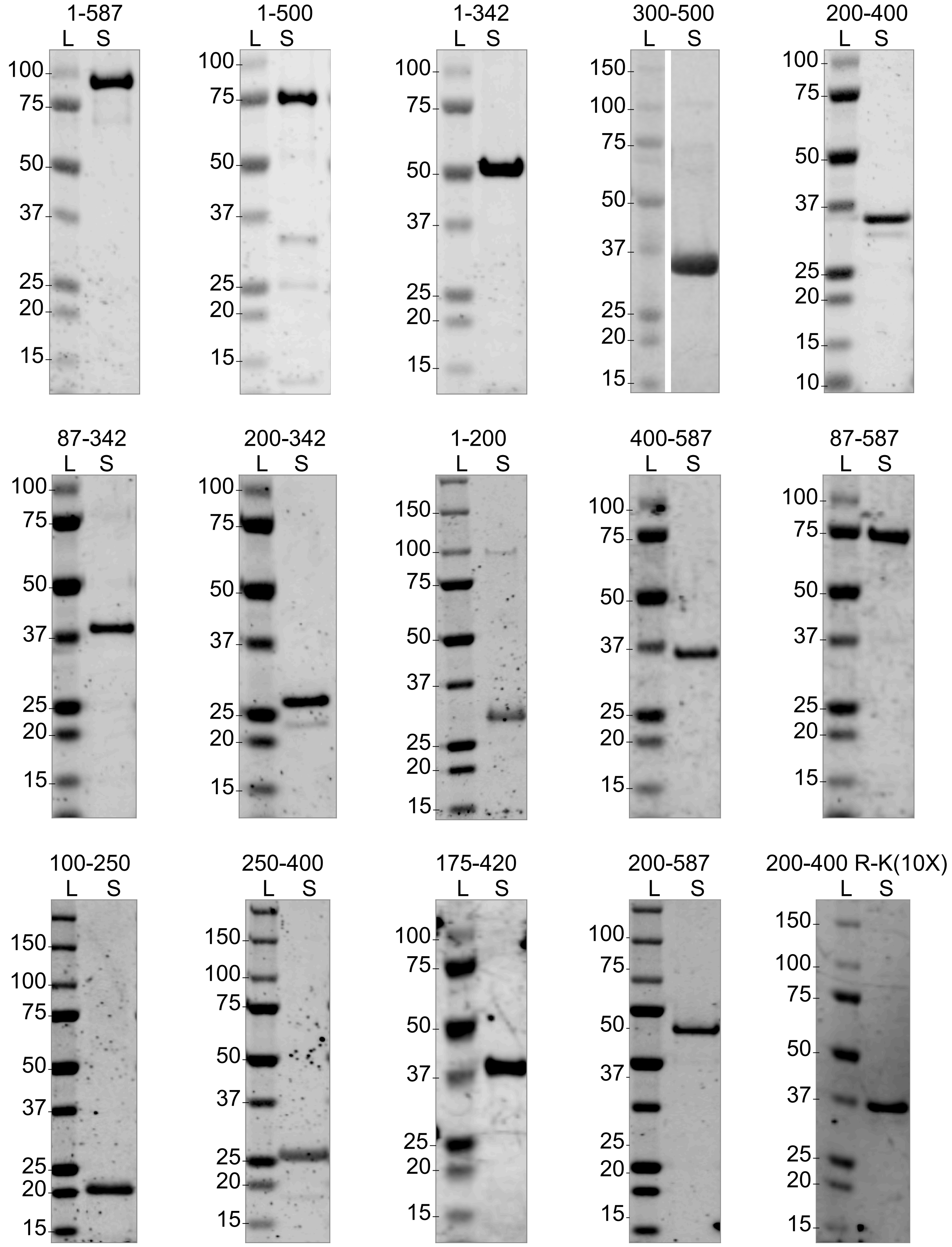

**Figure S1 (continued)**

**B** mNG-eIF5B-NTR

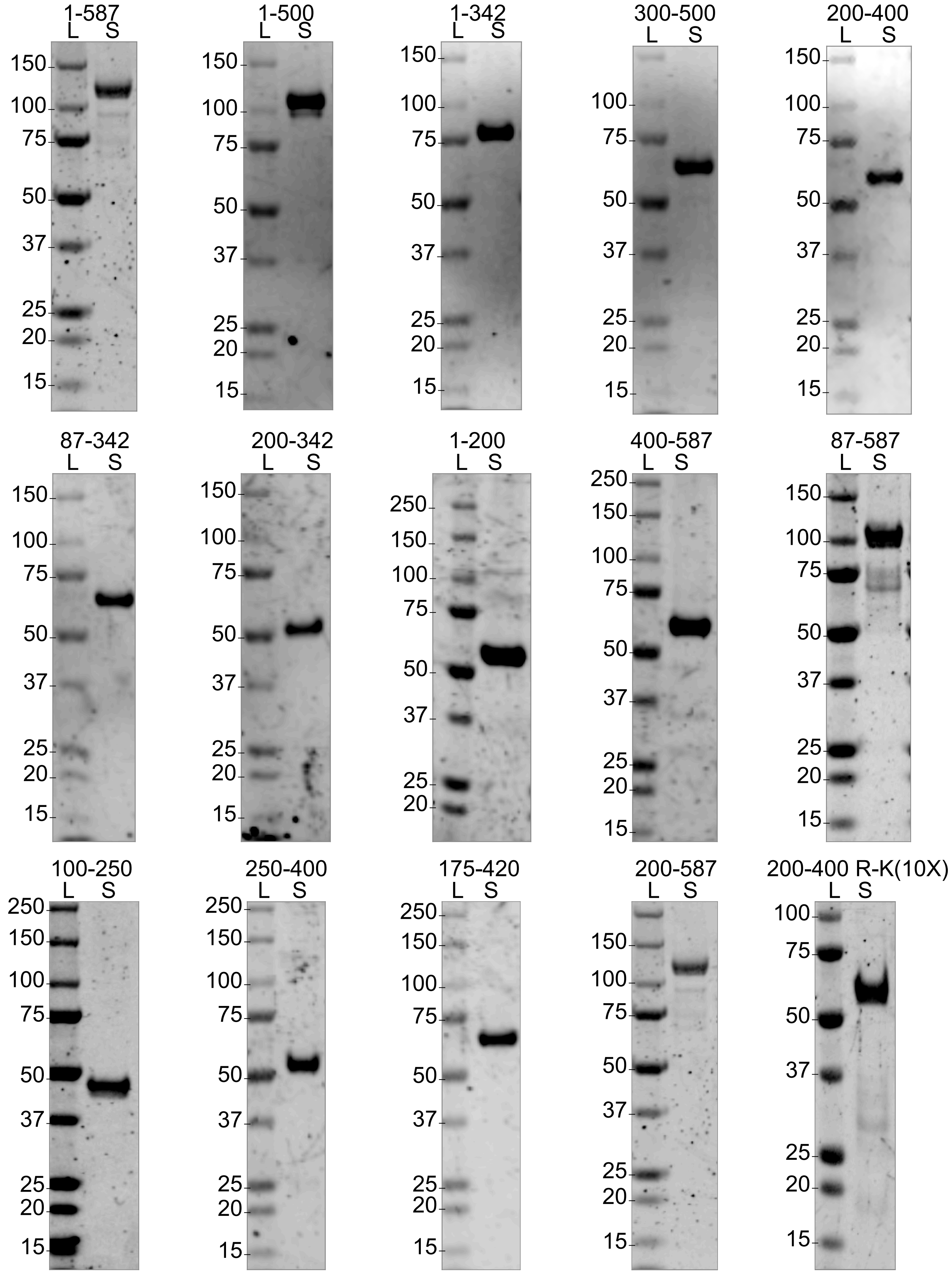

**Figure S1 (continued)**

**C**

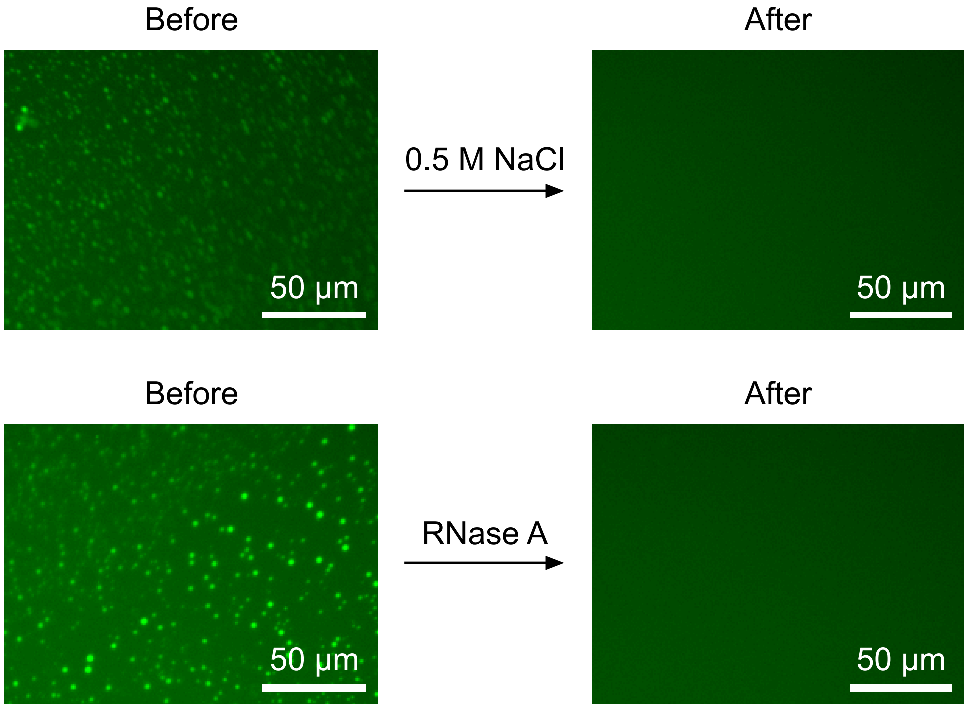


**Figure S1. Purified eIF5B-NTR proteins and reversibility of eIF5B-NTR droplets.** (**A**) SDS-PAGE of the purified untagged eIF5B-NTR constructs used in this study. (**B**) SDS-PAGE of the purified mNG-tagged eIF5B-NTR constructs used in this study. In (**A**) and (**B**), the purified protein sample (S) is shown beside the Precision Plus Protein standard (L) run on the same gel. Each panel is cropped to the region shown and adjusted only by linear contrast; molecular weights (kDa) are indicated on the left of each panel. The white vertical line in the 300–500 panel of (**A**) marks lanes from the same gel that were joined (spliced). (**C**) Widefield fluorescence images of eIF5B-NTR droplets formed with eIF5B-NTR (3.3 µM), with RNA (20 ng/µl) and 2% tracer (mNG-eIF5B-NTR) before and after addition of 0.5 M NaCl (top pair) or RNase A (bottom pair). Droplets dissolve completely under both conditions, indicating that condensation is RNA-dependent and electrostatically driven. Each image shows a 181 × 136 µm region of a 20× field; all four images are displayed with identical linear intensity scaling. Scale bars, 50 µm.

**Figure S2**

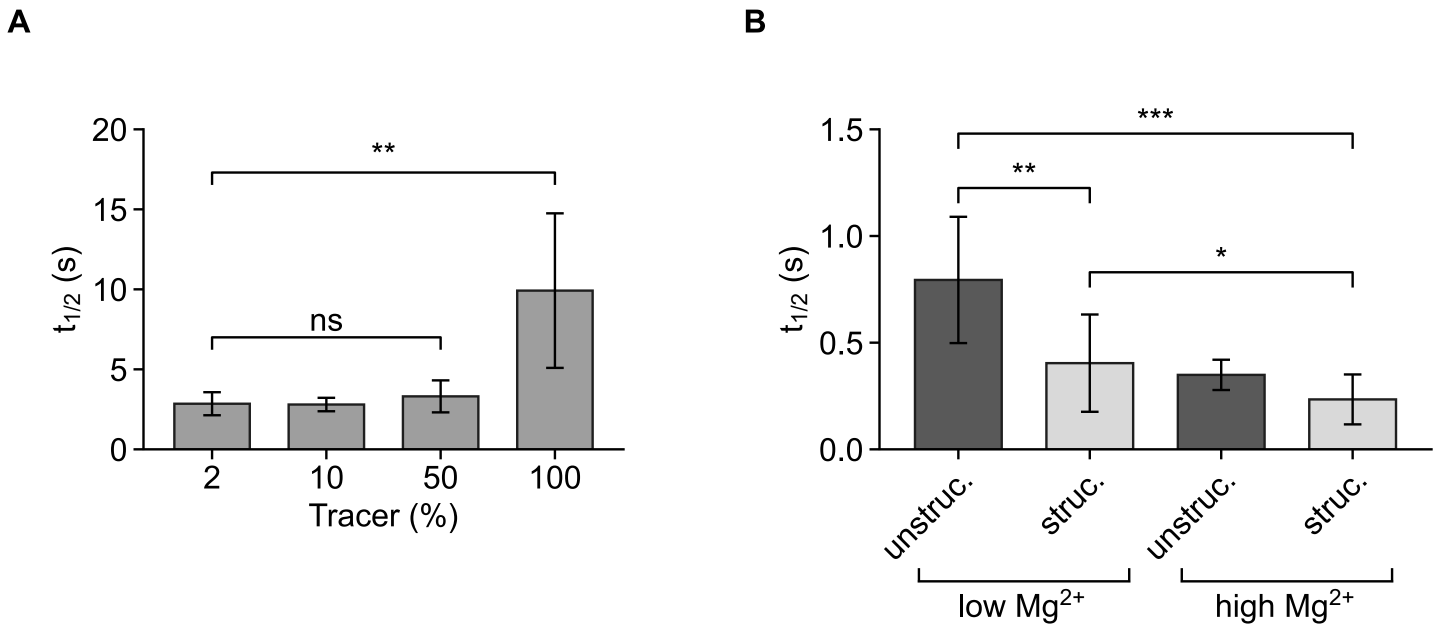


**Figure S2. Tracer controls and condensate dynamics with 1200-nucleotide RNA.** (**A**) FRAP recovery times ($t_{1/2}$) of eIF5B-NTR condensates formed with 2, 10, 50 or 100% mNG-eIF5B-NTR:eIF5B-NTR. Incorporation up to 50% does not alter condensate dynamics. Bars represent means ± SD of at least three independent replicates; unpaired t test; ns, $p \geq 0.05$; **, $p < 0.01$. (**B**) Recovery times of condensates formed with unstructured or structured 1200-nucleotide RNA at low (2 mM) or high (10 mM) $MgCl_2$. n = 15 droplets from three independent replicates. Bars represent mean ± SD; unpaired t test; *, $p < 0.05$; **, $p < 0.01$; ***, $p < 0.001$.

**Figure S3**

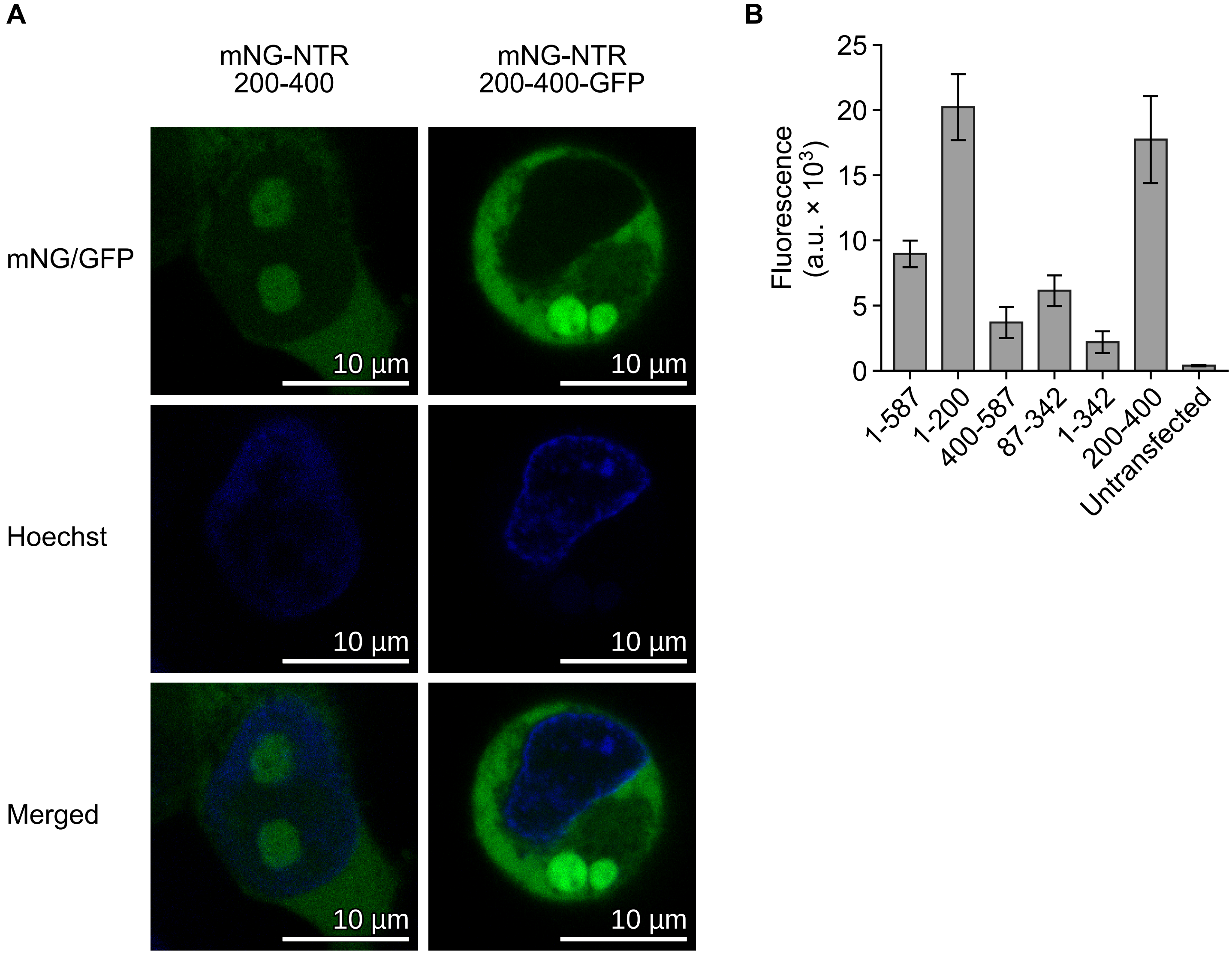


**Figure S3. Cellular expression and granule formation of eIF5B-NTR truncations.** (**A**) Representative confocal images of HEK293T cells expressing mNG-NTR$_{200–400}$ (left) or mNG-NTR$_{200–400}$-GFP (right) 48 h after transfection. The singly tagged core enters the nucleus and forms nuclear granules; addition of a second fluorescent tag (GFP) excludes it from the nucleus, so granules form in the cytoplasm. mNG/GFP (green), Hoechst-stained nuclei (blue) and merged images are shown without intensity adjustment. Scale bars, 10 µm. (**B**) Expression of mNG-tagged eIF5B-NTR truncations measured as total fluorescence of cell lysates 48 h after transfection and normalized to the number of fluorescent tags per construct; untransfected cells indicate background. Data represent mean ± SD of eight independent transfections.

**Table S1**

| Construct | Length (aa) | MW (kDa) | pI | Net charge (pH 7) | FCR | NCPR | κ |
|---|---|---|---|---|---|---|---|
| 1-587 | 593 | 68.11 | 4.97 | −36.8 | 0.55 | −0.06 | 0.24 |
| 1-500 | 509 | 59.18 | 6.01 | −2.9 | 0.53 | −0.01 | 0.20 |
| 1-342 | 341 | 38.66 | 8.79 | +6.1 | 0.53 | +0.02 | 0.23 |
| 87-342 | 256 | 29.25 | 9.18 | +10.0 | 0.55 | +0.04 | 0.23 |
| 300-500 | 217 | 25.39 | 5.47 | −5.0 | 0.56 | −0.02 | 0.16 |
| 200-342 | 148 | 16.97 | 9.60 | +13.0 | 0.56 | +0.09 | 0.16 |
| 200-400 | 204 | 24.20 | 9.41 | +12.0 | 0.62 | +0.06 | 0.11 |
| 1-200 | 201 | 22.66 | 5.38 | −4.0 | 0.51 | −0.02 | 0.29 |
| 400-587 | 195 | 22.21 | 4.21 | −43.9 | 0.51 | −0.23 | 0.35 |
| 175-420 | 249 | 29.28 | 9.48 | +15.0 | 0.59 | +0.06 | 0.12 |
| 200-587 | 395 | 45.86 | 4.89 | −31.8 | 0.57 | −0.08 | 0.23 |
| 87-587 | 508 | 58.70 | 4.97 | −32.8 | 0.56 | −0.07 | 0.25 |
| 250-400 | 154 | 18.39 | 9.15 | +6.0 | 0.64 | +0.04 | 0.09 |
| 100-250 | 154 | 17.56 | 8.77 | +3.0 | 0.51 | +0.02 | 0.30 |
| 200-400 R-K(10X) | 204 | 23.92 | 9.32 | +12.0 | 0.62 | +0.06 | 0.11 |

**Table S1 (continued)**

| Construct | Lys | Arg | K:R | Aromatic (%) | Predicted LLPS | Measured $C_{sat}$ (µM) |
|---|---|---|---|---|---|---|
| 1-587 | 120 | 24 | 5.00 | 1.9 | High | 1 |
| 1-500 | 112 | 24 | 4.67 | 2.1 | High | 1 |
| 1-342 | 85 | 9 | 9.44 | 2.6 | High | 0.5 |
| 87-342 | 67 | 8 | 8.38 | 3.1 | High | 2 |
| 300-500 | 43 | 15 | 2.87 | 0.9 | Medium | 10 |
| 200-342 | 42 | 6 | 7.00 | 1.4 | Low | 0.5 |
| 200-400 | 54 | 15 | 3.60 | 1.0 | Medium | 0.25 |
| 1-200 | 45 | 4 | 11.25 | 3.5 | Medium | n.d. |
| 400-587 | 22 | 6 | 3.67 | 1.0 | Low | n.d. |
| 175-420 | 60 | 21 | 2.86 | 1.2 | Medium | 0.25 |
| 200-587 | 76 | 20 | 3.80 | 1.0 | High | n.d. |
| 87-587 | 102 | 23 | 4.43 | 2.0 | High | 5 |
| 250-400 | 40 | 12 | 3.33 | 0.6 | Medium | 2 |
| 100-250 | 35 | 6 | 5.83 | 3.9 | Medium | 17 |
| 200-400 R-K(10X) | 64 | 5 | 12.80 | 1.0 | Low | 2 |

**Table S1. Physicochemical properties and phase-separation parameters of eIF5B-NTR constructs.** Predicted LLPS propensity was calculated with MolPhase (Liang *et al.* (2024) *EMBO J.* 43, 1898–1918). FCR, fraction of charged residues; NCPR, net charge per residue; κ, charge-patterning parameter; K:R, lysine-to-arginine ratio; Aromatic, percentage of aromatic residues; $C_{sat}$, measured saturation concentration (Figs. 3B and 4C); n.d., not determined (no condensates formed). 200-400 R-K(10X), core construct in which the ten arginines of patch 2 are substituted with lysine.
